# EGFR inhibition in lung adenocarcinoma upregulates cell surface expression of the placental antigen ALPP and enhances efficacy of ALPP-ADC therapy

**DOI:** 10.1101/2023.03.27.534173

**Authors:** Yihui Chen, Monica J. Hong, Hanwen Xu, Jody Vykoukal, Soyoung Park, Yining Cai, Ricardo A. León-Letelier, Ehsan Irajizad, Fu Chung Hsiao, Jennifer B. Dennison, Edwin J. Ostrin, Johannes F. Fahrmann, Hiroyuki Katayama, Samir M. Hanash

## Abstract

Alkaline phosphatase placental type (ALPP) and ALPPL2 are closely related and regulated GPI anchored proteins that are known to be expressed on the cell surface in some cancers, whereas normal tissue expression is largely limited to the placenta. Clinical utility of ALPP is potentially limited by heterogenous expression in tumors. Here, we assessed ALPP and ALPPL2 surfaceome protein levels in 158 cancer cell lines and mRNA expression levels in 10,967 tumors representing 32 cancer types from The Cancer Genome Atlas (TCGA), which revealed ALPP, and to a lesser extent ALPPL2, to be variably expressed in several cancer types including lung adenocarcinoma (LUAD). Surface expression of ALPP was confirmed by tissue microarray analysis of 204 lung tumors. Using LUAD as a model system, we demonstrated that treatment with EGFR inhibitors, or induction of cancer cell quiescence via nutrient deprivation greatly enhanced ALPP surface expression. Mechanistic studies revealed that enhancement of surface ALPP expression in LUAD following gefitinib treatment was mediated through repression of EGFR signaling and activation of the transcription factor FoxO3a, which was identified as an upstream transcriptional regulator of ALPP. Using xenograft models of LUAD, we further demonstrated that gefitinib treatment upregulates surface expression of ALPP in LUAD cells but not in normal tissues. Combination therapy with gefitinib and an ALPP antibody conjugated with Monomethylauristatin F (ALPP-ADC-MAF) resulted in superior anti-cancer efficacy compared with gefitinib or ALPP-ADC-MAF alone. Our findings support a novel combination treatment modality that boosts the efficacy of ALPP-ADC directed therapy.

## Introduction

Placental alkaline phosphatase (ALPP, also known as PLAP) and ALPPL2 (also known as ALPG) are members of the alkaline phosphatase family that are exclusively expressed in the placenta and share 98% similarity in amino acid sequence [1, 2]. Several studies have documented heterogenous expression of ALPP and ALPPL2 in various cancer types [1, 2, 3, 4, 5, 6]. The restricted expression of ALPP and ALPPL2 in normal tissues and their accessibility on the cancer cell surface supports their potential as targets for cancer therapeutics. To this end, a chemical library screen resulted in a selective and potent ALPP inhibitor which specifically bound to ALPP-positive tumors *in vitro* and targeted cervical cancer in a mouse model of the disease [4]. A fluorescent derivative of the ALPP inhibitor functioned as a bispecific engager directing chimeric antigen receptor-T cells to fluorescein on ALPP-positive tumor cells for chimeric antigen receptor (CAR) T-cell mediated cancer killing [4]. Targeting ALPP with a α-ALPP and α-CD3 bispecific antibody showed specific killing of ALPP-positive colorectal cancer cell lines [7]. Expression of ALPP in colorectal cancer led to its consideration for CAR T-cell therapy [5]. ALPP-CAR-T cells mediated potent cytotoxicity towards cancer cells, while the combination of ALPP-CAR-T cells with anti-PD-1, PD-L1, or LAG-3 checkpoint inhibitors further increased the therapeutic efficacy of CAR T-cells [5]. Second-generation CAR T-cells with a fully human scFv against ALPP antigen effectively killed ALPP-expressing HeLa cells [8]. A clinical trial using α-ALPP CAR T cells for ovarian and endometrial cancer has been initiated, while another clinical study evaluating the efficacy of ALPP/ALPPL2 antibody-drug conjugate in advanced solid tumors is ongoing [9].

Yet, the expression levels of ALPP and ALPPL2 in most cancer types are relatively low. Assessment of ALPP protein levels in 12,381 tumors by immunohistochemistry showed strong expression of ALPP in more than 50% of seminoma, embryonal carcinoma of the testis, and yolk sac tumor, which dropped to less than 20% in all other cancer types examined [6]. Nevertheless, the ratio of ALPP positive tumors are significantly higher with 6.1% weak, 1.5% moderate, and 4.5% strong staining of ALPP in 12,281 tumor specimens [6]. For lung cancers, ALPP is positive in 20.6% of adenocarcinoma, 1.5% of squamous cell carcinoma, and 0% in small cell carcinoma, with more than 95% with weak expression [6]. Thus, few patients may benefit from the ALPP targeting therapies, but upregulating ALPP expression may be an attractive strategy to broaden the application.

In the current study, we evaluated mRNA expression patterns of ALPP and ALPPL2 in 10,967 tumors representing 32 cancer types from The Cancer Genome Atlas (TCGA) as well as surfaceome protein levels in 158 cancer cell lines. Analyses revealed ALPP, and to a lesser extent ALPPL2, to be variably expressed in several cancer types, including high expressing outliers in lung adenocarcinoma (LUAD), which was further independently confirmed in a tissue microarray of LUAD tumors. Using LUAD as a model system, we report a novel finding that induction of cancer cell quiescence via nutrient deprivation, cell-cell contact inhibition as well as treatment with chemotherapeutic or targeted agents, including the EGFR-targeting inhibitor gefitinib, amplifies ALPP surface expression on cancer cells. Mechanistic studies revealed FoxO3a as the upstream transcriptional regulator of ALPP. Combination therapy with gefitinib and an ALPP antibody conjugated with monomethylauristatin F (ALPP-ADC-MAF) resulted in enhanced anti-cancer effects compared with gefitinib or ALPP-ADC-MAF alone in vitro and in a xenograft mouse model of LUAD.

## Materials and Methods

### Cell culture

Detailed information on the human cancer cell lines used in this study is provided in **Supplementary Table S1**. Cells were culture in Roswell Park Memorial Institute 1640 Medium (RPMI 1640, Cat. #10-040-CV, Corning) supplemented with 10% inactivated fetal bovine serum (FBS, Cat. #16140-071, Gibco) and maintained at 37 °C in a humidified atmosphere with 5% CO_2_.

### Antibodies, chemicals, and virus strains

Detailed information on the antibodies, chemicals, and virus strains used in this study is provided in **Supplementary Table S2**. ALPP antibody (Cat. #NB110-3638, Novus biologicals) was conjugated with Monomethyl auristatin F (MMAF) by the Creative Biolabs.

### Proteomics Analysis

Proteomics analysis was performed as previously described [10, 11, 12, 13, 14]. For proteomic analysis of whole-cell extract (WCE), 2 × 10^7^ cells were lysed in 1 mL of Tris-HCl (pH 8.0) containing 3% octyl-glucoside (1% w/v), 4M urea, and protease inhibitors (complete protease inhibitor cocktail, RocheDiagnostics), followed by sonication and centrifugation at 20,000g with collection of the supernatant, and filtration through a 0.22µm filter. For surfaceome, 4 × 10^8^ cells were labeled with EZ-link Sulfo-NHS-SS-Biotin reagent (0.8mM, Cat. #21331, Thermo Scientific) for 30 minutes at room temperature, followed by quenching of reaction with lysine solution (10 mM) for 15 minutes at room temperature. Cells were then washed with PBS for 3 times and lysed in Tris-HCl (pH 8.0) containing 2% octyl-glucoside. After sonication and centrifugation at 20,000g, the collected supernatant was incubated NeutrAvidin Plus UltraLink Resin (Cat. #53151, Thermo Scientific) for 4 hours at 4°C, followed thrice PBS washes, and subsequent incubation with PBS containing Tris-HCl (100 mM, pH8.0) and DTT (1% w/v) overnight at 4°C. After centrifugation, biotinylated surface proteins were collected for further process.

Two milligrams of WCE or surface proteins were reduced in DTT and alkylated with acrylamide before fractionation with RP-HPLC. A total of 84 fractions were collected at a rate of 3 fractions/min. Mobile phase A consisted of H2O:ACN (95:5, v/v) with 0.1% of TFA. Mobile phase B consisted of ACN:H2O (95:5) with 0.1% of TFA. Collected fractions from were dried by lyophilization followed by in-solution digestion with trypsin (Mass Spectrometry Grade, Thermo Fisher).

Based on the chromatogram profile, 84 fractions were pooled into 24 fractions for LC-MS/MS analysis per cell line. In total, 2,688 fractions were analyzed by RPLC-MS/MS using a nanoflow LC system (EASYnano HPLC system, Thermo Scientific) coupled online with LTQ Orbitrap ELITE mass spectrometer (Thermo Scientific). Separations were performed using 75 µm id × 360 µm od × 25-cm-long fused-silica capillary column (Column Technology) slurry packed with 3 µm, 100 A° pore size C18 silica-bonded stationary phase. Following injection of 500 ng of protein digest onto a C18 trap column (Waters, 180 µm id×20 mm), peptides were eluted using a linear gradient of 0.35% mobile phase B (0.1 formic acid in ACN) per minute for 90 min, then to 95% B in for 10 min at a flow rate of 300 nL/min. Eluted peptides were analyzed by LTQ Orbitrap ELITE in data-dependent acquisition mode. Each full MS scan (m/z 400–1800) was followed by 20 MS/MS scans (CID normalized collision energy of 35%). Acquisition of each full mass spectrum was followed by the acquisition of MS/MS spectra for the 20 most intense +2, +3 or +4 ions within a duty cycle. Dynamic exclusion was enabled to minimize redundant selection of peptides previously selected for MS/MS analysis. Parameters for MS1 were 60,000 for resolution, 1 × 106 for automatic gain control target, and 150 ms for maximum injection time. MS/MS was done by CID fragmentation with 3×10^4^ for automatic gain control, 10 ms for maximum injection time, 35 for normalized collision energy, 2.0 m/z for isolation width, 0.25 for activation q-value, and 10 ms for activation time.

MS/MS spectra were searched against the Uniprot proteome database using X!Tandem in Trans-Proteomic pipeline (TPP-ver4.8). One fixed modification of carbamidomethyl at Cys (57.02146 Da) Propionamide at Cys (71.037114 Da) and variable modifications, oxidation at Met (15.9949 Da) and SILAC ^13^C6 at Lys (6.0201 Da) were chosen. The purpose of adding SILAC ^13^C6 was to filter out the possible Bovine proteins contaminated from the culturing media. The mass error allowed was 20 ppm for parent monoisotopic and 0.5 Da for MS2 fragment monoisotopic ions. Full trypsin was specified as protein cleavage site, with possibility of two missed cleavages allowed. The searched result was filtered with FDR=0.01.

Each dataset was normalized to the total number of spectral counts of the each compartment [15].

### FoxO3a overexpression

The open reading frame of FoxO3a was inserted into pLOC-RFP vector and packed into lentivirus. Stable FoxO3a overexpression in LUAD cell lines HCC827 and H1650 was achieved by lentivirus infection and subsequent selection of blasticidin-resistant cells and overexpression of FoxO3a was verified by immunoblots.

### Immunofluorescence

A total of 50,000 cells were plated on coverslips, fixed with 4% paraformaldehyde, permeabilized with 0.02 Triton X-100, and blocked with 5% donkey serum. Cells were subsequently incubated with α-ALPP antibody (Cat. #NB110-3638, Novus biologicals) diluted 1:100 in PBS containing 0.1% Bovine serum albumin overnight at 4°C, and then secondary antibody for 1 hour in the dark at room temperature. Images were acquired in z-series on a spinning-disk confocal system.

### Hematoxylin and Eosin (H&E) staining

H&E staining was performed following the manufacturer’s instructions (Cat. #ab245880, Abcam). The tissue sections were deparaffinized and hydrated by 3 washes in Xylene and serial washes in 100%, 95%, 70%, and 50% ethanol, followed by 2 washes in deionized water. Hematoxylin, Mayer’s (Lillie’s Modification) solution was applied to tissue sections and incubated for 5 minutes. Tissue slides were rinsed in distilled water and stained with Bluing Reagent for 15 seconds. After another 2 washes with distilled water, the slides were dipped into 100% ethanol. Subsequently, the slides were stained with Eosin Y Solution (Modified Alcoholic) for 3 minutes, and dehydrated in 100% ethanol, followed by imaging under an inverted microscope.

### Immunohistochemistry (IHC) analysis

Tissue microarray for IHC staining of ALPP in this study comprised of 204 surgically resected lung cancer tumor specimens collected under an institutional review board protocol and archived as formalin-fixed, paraffin-embedded specimens in The University of Texas Specialized Program of Research Excellence thoracic tissue bank at The University of Texas MD Anderson Cancer Center. Patient characteristics for the analyzed cohort are provided in Supplementary Table S3. Human normal tissue microarray for IHC staining of ALPP was obtained from Novus Biologicals (Cat. #NBP2-78113). IHC staining was performed as previously described [16]. Briefly, IHC staining of ALPP (Cat. #ab133602, Abcam) was performed on 5 µm of unstained sections from tissue microarray and mouse tissue blocks constructed using formalin-fixed paraffin-embedded tissues.

### Cell proliferation assay

The LUAD cell lines HCC827 and H1650 were seeded in a 96-well plate at a density of 1 × 10^5^ cells/well. Cell viability was determined using the CellTiter 96 Aqueous One Solution Cell Proliferation Assay (MTS) kit (Cat. #G3580, Promega) at 24, 48, and 72 hours post-treatment. All experiments were performed in biological triplicates.

### Chromatin immunoprecipitation (ChIP)

ChIP was performed in HCC827 and H1650 cells using Pierce Magnetic ChIP Kit (Cat. #26157, Thermo Scientific) following the manufacturer’s instruction. Briefly, cells were seeded in a 100 mm petri dish at a density of 80% confluency and treated with DMSO or gefitinib (1 µM) for 6 hours; 2 x 10^6^ cells were subsequently used for each ChIP assay. After DMSO or gefitinib treatment, crosslinking was performed in cell culture medium with 1% formaldehyde for 10 minutes at room temperature, followed by addition of glycine solution and incubation for 5 minutes. Cells were washed twice with ice-cold PBS and scraped off in 1 mL ice-cold PBS with Halt Protease Inhibitor Cocktail. Cell pellets obtained after centrifugation at 3,000 x g for 5 minutes were resuspended in 200 µL Membrane Extraction Buffer, vortexed for 15 seconds, and incubated on ice for 10 minutes. Following centrifugation at 9,000 x g for 3 minutes, nuclei pellets were resuspended in 200 µL MNase Digestion Buffer Working Solution with MNase (2 Units) and incubated in a 37°C water bath for 15 minutes with periodic mixing. The DNA digestion reaction was terminated by the addition of 20 µL MNase Stop Solution and incubation on ice for 5 minutes. Nuclei were spun down and resuspended in 100 µL IP Dilution Buffer, followed by sonication with 20-second pluses/20-second on ice for three sequential cycles and centrifugation. Thereafter, 90 µL of supernatant was mixed with 410 µL of IP Dilution Buffer and antibodies (Positive antibody: 10 µL; Negative antibody: 2 µL; FoxO3a antibody: 8 µL (Cat. #720128, Invitrogen)) and incubated overnight at 4°C. Subsequently, the DNA-protein complexes were enriched with 20 µL of Protein A/G magnetic Beads and eluted in IP Elution Buffer. Proteins were digested with addition of proteinase K and DNA recovered by the DNA Clean-up column. The relative abundance of genomic DNA fragments was determined using Sso Advanced Universal SYBR Green Supermix (Cat. #1725271, Bio-Rad) with the following primers specific for the promoter region of human *ALPP* gene.

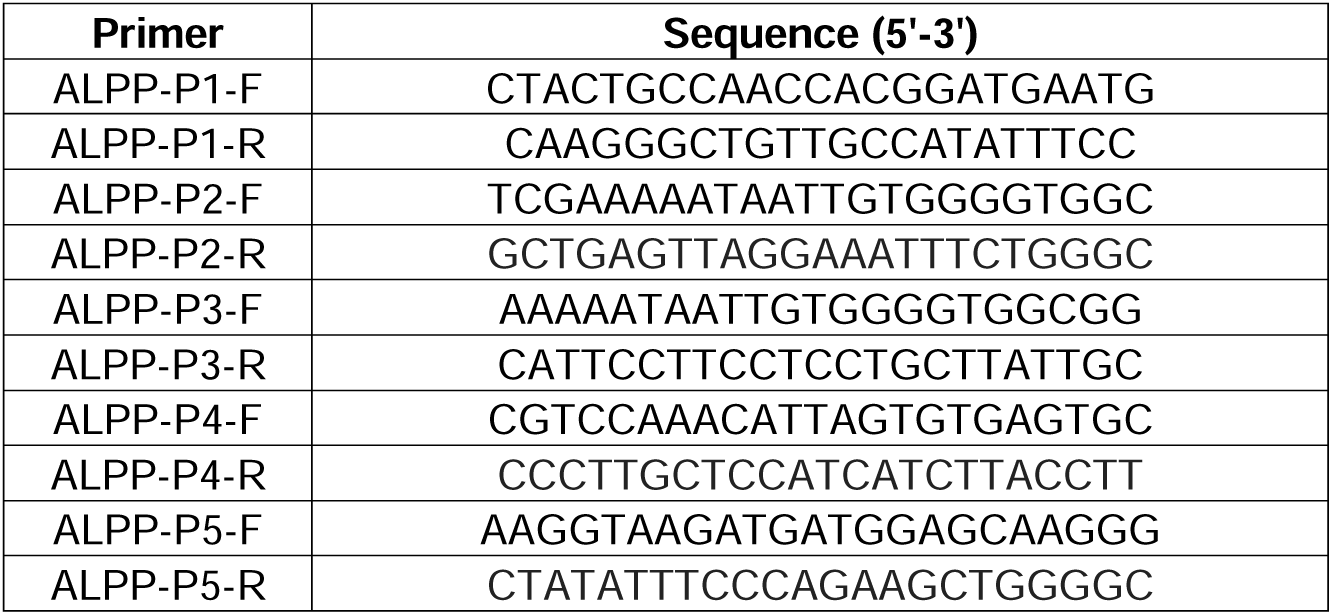

### Total RNA extraction and quantitative PCR (qPCR)

Total RNA extraction and qPCR was performed as previously described [13, 17]. Briefly, Total RNA was extracted using the RNeasy Mini Kit (Cat. #74104, QIAGEN). Reverse transcription of total RNA was performed using the High-Capacity Reverse Transcriptase Kit (Cat. #4368813, Applied Biosystems). TaqMan Universal Master Mix II (Cat. #4440040, Applied Biosystems) was used for analyzing ALPP expression. Probe for ALPP was purchased from Thermo Fisher Scientific (Hs03046558_s1).

### Immunoblotting

Total proteins were extracted in RIPA buffer (Cat. #89901, Pierce) containing complete protease inhibitor cocktails (Cat. #04693116001, Roche) and phosSTOP (Cat. #04906837001, Roche) and resolved by 4-15% gradient SDS-PAGE (Cat. #64503191, #64485082, BioRad) at 110 V, and subsequently transferred to PVDF membranes using the Trans-Blot Turbo system (BioRad). Membranes were blocked with 5% non-fat Blotting-Grade Blocker (Cat. #1706404, BioRad) for 1 hour at room temperature and incubated with primary antibodies at 4 °C overnight. Information for antibodies is provided in Supplementary Table S1. After three washes with Tris-buffered saline with 0.1% Tween® 20 Detergent (TBST), membranes were incubated with appropriate HRP-conjugated secondary antibodies for 1 hour at room temperature, followed by another three washes with TBST. Membranes were visualized using ECL2 reagent substrate (Cat. #170-5061, BioRad).

### Cell cycle analysis

The LUAD cell lines HCC827 and H1650 were harvested after indicated treatment for flow cytometry-based cell cycle analysis. Briefly, cells were washed three times with ice-cold phosphate-buffered saline (PBS) and fixed in 70% ethanol at −20°C overnight. After another three PBS washes, cells were resuspended and stained in 0.5 mL of propidium iodide staining solution (PI, Cat. #00-6990-50, Invitrogen, MA, USA) for 30 minutes in the dark at room temperature, followed by cell cycle analyses using a Gallios Flow Cytometry System.

### Antibody internalization assay

Antibody internalization was assessed using the pHrodo™ iFL Red Microscale Protein Labeling Kit (Cat. #P36014, Thermo Fisher) following the manufacturer’s instructions. Briefly, 100 µg of anti-ALPP antibody (Cat. #MAB59051, R&D systems) in 100 µl PBS was mixed with 10µl of 1 M sodium bicarbonate and 5 µl of 2 mM pHrodo followed by incubation for 30 minutes at room temperature. The pHrodo-conjugated anti-ALPP antibody was subsequently purified using the gel resin. Cells were washed twice with PBS and incubated with pHrodo-conjugated anti-ALPP antibody (10 µg/ml) for 24 hours, followed by fixation in 2% paraformaldehyde, two washes with PBS, and mounting with DAPI. Images were captured using a fluorescent microscope.

### Animal study

Animal experiment protocols were approved by The University of Texas MD Anderson Cancer Center IRB and in accordance with the Guidelines for the Care and Use of Laboratory Animals published by the NIH (Bethesda, MD). BALB/c nude mice (Cat. #194, Charles River) were housed in specific pathogen free facilities. For orthotopic xenograft models of LUAD, a total of 1 x 10^6^ cells H1650 cell lines expressing firefly luciferase were suspended in 50 µl 50% Matrigel Matrix (Corning)/Opti-MEM media and injected into the lung of 8- to 10-week-old nude mice. For gefitinib treatment, mice received gefitinib (50mg/kg) via oral gavage daily for 10 days starting from Day 17 post cancer cell inoculation. For the gefitinib/ALPP-ADC combinational treatment, mice received additional ALPP antibody conjugated with monomethylauristatin F (anti-ALPP-MMAF) (5mg/kg, i.v.) at Day 19 and Day 26. The tumor growth was monitored twice a week over a period of 5 weeks using the Xenogeny In Vivo Imaging System (IVIS, Alameda, CA). Mice were euthanized at Day 30 and the tumors were harvested and processed for routine histological and immunohistochemical analyses.

### Gene expression datasets

Gene expression for ALPP and ALPPL2 for The Cancer Genome Atlas (TCGA) PanCancer Atlas Studies representing 10,967 tumors across 32 cancer types was downloaded from cbioportal (https://www.cbioportal.org/).[18] Gene expression for ALPP and ALPG (ALPPL2) in 1,037 cancer cell lines in the Cancer Cell Line Encyclopedia (CCLE) was downloaded from the Broad Institute (https://sites.broadinstitute.org/ccle/).

### Statistical analysis

For continuous variables, statistical significance was determined by 2-sided Student T-Test unless otherwise specified. For categorical variables, statistical significance was determined by Fisher’s Exact Test for two class categorical comparisons or χ^2^ test for trend for multiple categorical variables. Figures were generated in GraphPad Prism Version 9.0.0.

## Results

### ALPP and ALPPL2 are expressed on the cell surface of a variety of cancer types

Assessment of ALPP and ALPPL2 in the whole cell extract (WCE) and the surface compartment of 158 cancer cell lines similarly revealed ALPP, and to a lesser extent ALPPL2, to be variably expressed in several cancer types, with 34 (21.5%) and 48 (30.4%) of analyzed cell lines having detectable levels ALPP in the WCE or surface compartment. Surface ALPP expression was particularly enriched in breast, ovarian, colon, pancreatic, gastric, and LUAD cancer cells (**Figure 1A**). ALPP and ALPPL2 tended to be co-expressed on the surface of cancer cells (Spearman Rho: 0.69 (95% CI: 0.59-0.76)).

**Figure 1.**
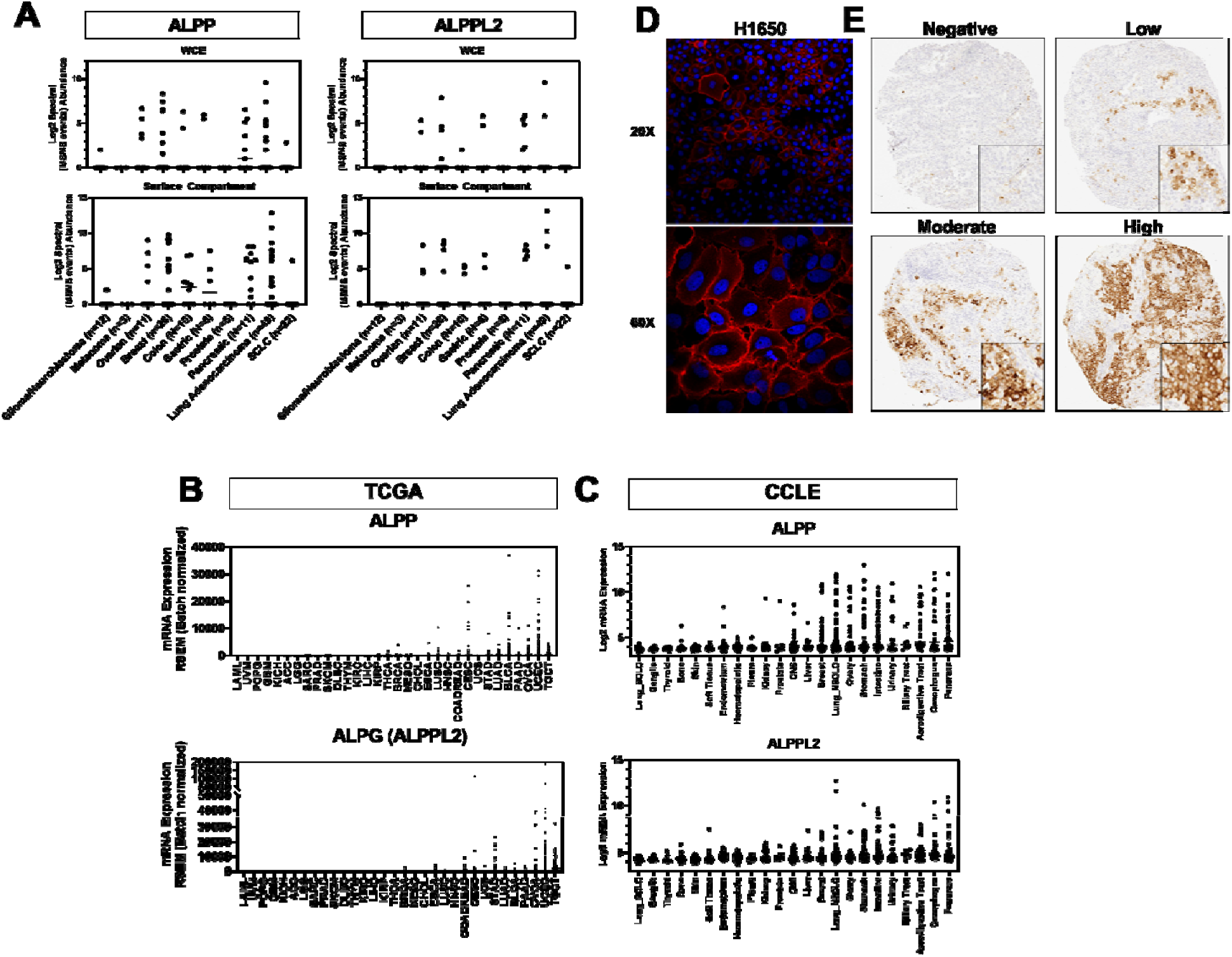
Surface ALPP and ALPPL2 expression in different cancer types. **A)** Log2 spectral abundance (MSMS events) of ALPP and ALPP2 in whole cell extracts (WCE) and the surface compartment of 158 cancer cell lines. Numbers in parentheses indicates the number of cell lines analyzed for a given cancer type. **B)** Gene expression levels of ALPP and ALG (ALPPL2) in 10,967 tumors representing 32 cancer types in The Cancer Genome Atlas (TCGA) datasets. Cancer types are ranked by average ALPP mRNA expression from lowest to highest. Abbreviations: **ACC**: Adrenocortical carcinoma; **BLCA**: Bladder Urothelial Carcinoma; **BRCA**: Breast invasive carcinoma; **CES**: Cervical squamous cell carcinoma and endocervical adenocarcinoma; **CHOL**: Cholangiocarcinoma; **COAD**: Colon adenocarcinoma; **DLBC**: Lymphoid Neoplasm Diffuse Large B-cell Lymphoma; **ESCA**: Esophageal carcinoma; **GBM**; Glioblastoma multiforme; **HNSC**; Head and Neck squamous cell carcinoma; **KICH**: Kidney Chromophobe; **KIRC**: Kidney renal clear cell carcinoma; **KIRP**: Kidney renal papillary cell carcinoma; **LAML**: Acute Myeloid Leukemia; **LCML**: Chronic Myelogenous Leukemia; **LGG**: Brain Lower Grade Glioma; **LIHC**: Liver hepatocellular carcinoma; **LUAD**: Lung adenocarcinoma; **LUSC**: Lung squamous cell carcinoma; **MESO**: Mesothelioma; OV: Ovarian serous cystadenocarcinoma; **PAAD**: Pancreatic adenocarcinoma; **PCPG**: Pheochromocytoma and Paraganglioma; **PRAD**: Prostate adenocarcinoma; **READ**: Rectum adenocarcinoma; **SARC**: Sarcoma; **SKCM**: Skin Cutaneous Melanoma; **STAD**: Stomach adenocarcinoma; **TGCT**: Testicular Germ Cell Tumors; **THCA**: Thyroid carcinoma; **THYM**: Thymoma; **UCEC**: Uterine Corpus Endometrial Carcinoma; **UCS**: Uterine Carcinosarcoma; **UVM**: Uveal Melanoma. **C)** Gene expression of ALPP and ALPPL2 in 1,037 cancer cells stratified by anatomical site from the Cancer Cell Line Encyclopedia (CCLE). Cancer types are ranked by averaged ALPP mRNA expression from lowest to highest. **D)** Immunofluorescence staining of ALPP in LUAD *EGFR-mutant* H1650 cell line. **E)** Representative IHC sections for negative-, low-, moderate-, and high-ALPP membrane staining in lung tumors.

We additionally evaluated mRNA expression levels of ALPP and its paralogue ALPPL2 (encoded by the *ALPG* gene) in 10,967 tumors representing 32 cancer types from The Cancer Genome Atlas (TCGA) and 1,037 cancer cell lines from the Cancer Cell Line Encyclopedia (CCLE) which revealed heterogenous expression patterns with gastrointestinal and gynecological cancers as well as in lung adenocarcinoma (LUAD) exhibiting evident high-expressing outliers (**Figure 1B-C**).

Focusing on ALPP, we first confirmed surface expression via immunofluorescence in *EGFR*mutant H1650 LUAD cells, which had high surface ALPP expression based on mass spectrometry findings (**Figure 1D**). Using a tissue microarray (TMA) consisting of 140 LUAD and 64 squamous cell carcinoma (SCC) tumors, we further assessed ALPP expression by immunohistochemistry (IHC). Of the 204 tumors, 37 (18.1%) stained positive for cytosolic ALPP whereas 23 (11.3%) stained positive for membranous ALPP. Representative IHC staining for negative-, low-, moderate-, and high-ALPP membrane staining in lung tumors is provided in **Figure 1E**. ALPP membrane staining positivity was more frequent in LUAD tumors (21 of 119) than SCC tumors (2 out of 64; Fisher’s exact test p-value: 0.015), consistent with prior reports [6]. ALPP membrane staining positivity associated with smoking status (χ^2^ test for trend p-value: 0.0001) (**Supplemental Table S3**). No statistically significant associations were found between tumoral ALPP membrane staining positivity and sex, stage, or mutational status (**Supplemental Table S1**). Except for placental and testis tissues, ALPP protein staining in other normal tissues, including lung, were negative (**Supplemental Figure S1A-B**).

### ALPP expression is potentiated by anti-proliferative agents

Previous studies have reported an inverse association between ALPP expression and cell proliferation [19, 20, 21, 22, 23], although the underlying mechanism(s) remain largely unknown. To test this phenomena, we inhibited cell proliferation in *EGFR-*mutant HCC827 and H1650 LUAD cells via FBS-restriction or high-confluency contact inhibition [24] and assessed for changes in ALPP protein expression. Both FBS-restriction and high-confluency contact inhibition led to marked induction of ALPP protein expression in HCC827 and H1650 cell lines (**Figures 2A-B**). We further treated H1650 LUAD cells with the anti-proliferative chemotherapeutic agents 5-fluorouracil (5-FU) or gemcitabine and similarly found pronounced increases in ALPP mRNA and protein expression compared to vehicle control (**Figure 2C and supplementary Figure S2**).

**Figure 2.**
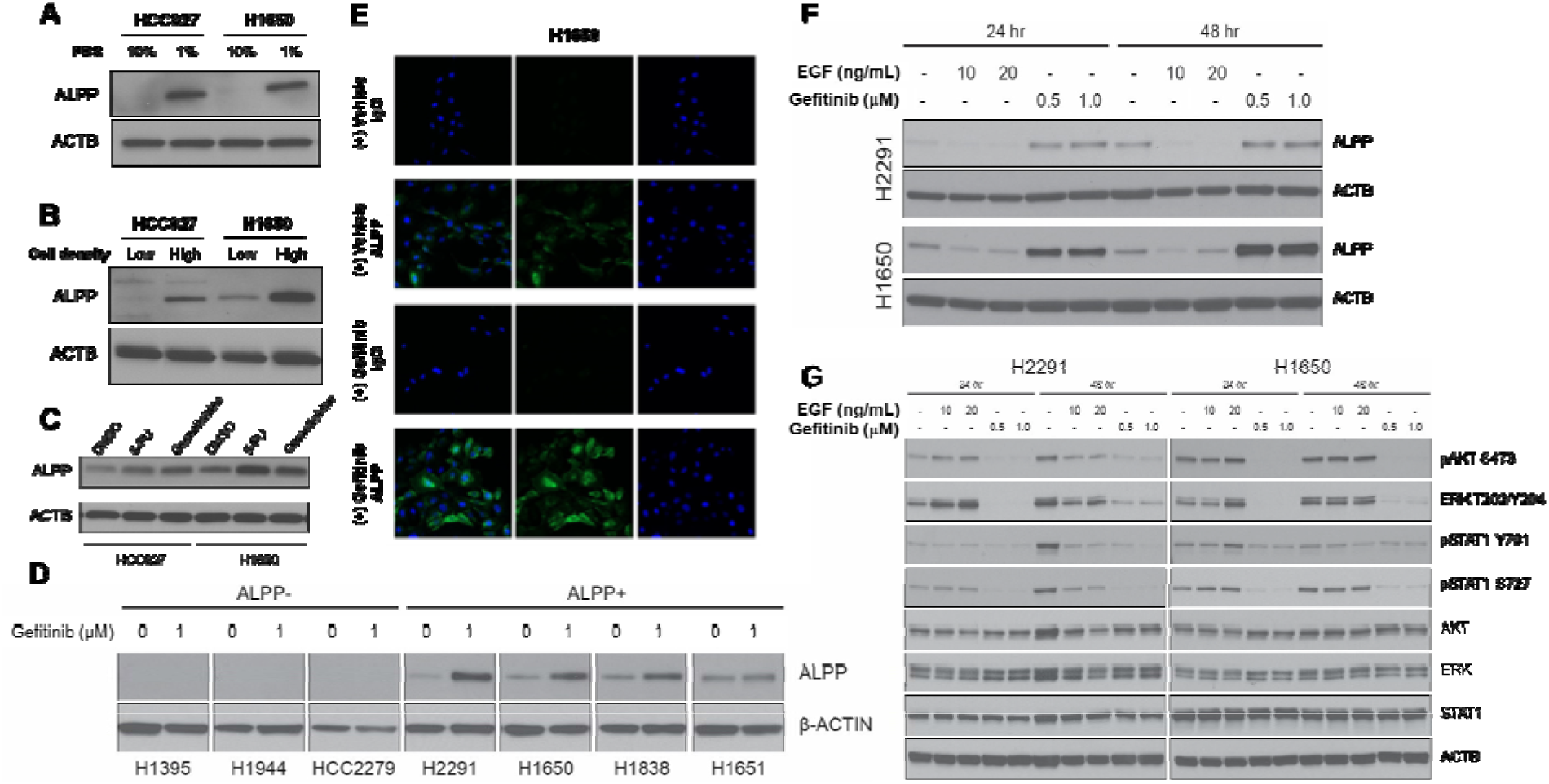
Growth inhibition potentiates ALPP expression. (A) Immunoblots for ALPP in HC827 and H1650 LUAD cells cultured in RPMI 1640 medium supplemented with 10% or 1% FBS for 48 hours. (B) Immunoblots for ALPP in HC827 and H1650 LUAD cells at high (80%) or low (30%) cell density. (C) Immunoblots for ALPP in HC827 and H1650 LUAD cells following 48-hour treatment with either vehicle, 5-FU (10 µM), gemcitabine (1 µM), or paclitaxel (10 nM). (D) Immunoblots for ALPP in LUAD cells with basal ALPP expression (H2291, H1650, H1838, and H1651) and LUAD cells (H1395, H1944, HCC2279) that did not express ALPP at the basal level. Cells were treated with either vehicle or gefitinib (1 µM) for 48 hours. (E) Immunofluorescence staining of ALPP using IgG isotope control or α-ALPP antibody in H1650 LUAD cells following 24-hour treatment with either vehicle (DMSO) or gefitinib (1µM). (F-G) Immunoblots for ALPP, pAKT-S473, total AKT, ERK-T202/Y204, total ERK, pSTAT1-Y701, pSTAT1-S727, total STAT1, and D-actin (loading control) in LUAD cancer cell H2291 and H1650 following 24- and 48-hour treatment with or without EGF (10ng/mL) or gefitinib (1µM).

Tyrosine kinase inhibitors (TKIs) are frequently used anti-proliferative agents for various cancers, including *EGFR-*mutant LUAD.[25, 26] Treatment of *EGFR-*mutant H1650 LUAD cells with the EGFR inhibitor gefitinib resulted in increased ALPP protein levels and enhanced ALPP surface expression (**Figure 2D-E**). Treatment of H1650 and HCC827 LUAD cells with other TKIs including lapatinib, afatinib, and osimertinib yielded similar results (**Supplemental Figure S3**). TKI-mediated enhancement of ALPP expression was also observed in *EGFR* wild-type (wt) H2291, H1838, and H1651 LUAD cell lines as well as in breast cancer (BT20 and HCC1937) and pancreatic cancer (BxPC-3, PANC03.27, SW1990, AsPC-1, and SU.86.86) cell lines, indicating that this effect is not restricted to *EGFR*-mutant LUAD (**Figure 2D; Supplemental Figure S4A-B**).

To test whether enhanced ALPP expression was mediated through EGFR signaling inhibition, we cultured *EGFR*-mutant H1650 and *EGFR*-wt H2291 LUAD cells in growth medium with or without epidermal growth factor (EGF), the endogenous ligand of EGFR. ALPP expression was drastically reduced in both cell lines cultured in EGF-containing growth medium (**Figure 2F**). Supplementation of H1650 and H2291 LUAD cells with EGF induced AKT phosphorylation at serine 473 (Ser473), and ERK phosphorylation at threonine 202 (T202) and tyrosine 204 (Y204) residues, whereas gefitinib-treatment repressed phosphorylation of AKT and ERK (**Figure 2G**).

### ALPP is transcriptionally regulated by FoxO3a

FoxO family transcription factors are involved in cell cycle regulation and are regulated by PI3K/AKT and MEK/ERK signaling [27]. At the mRNA level, *EGFR*-mutant H1650 and HCC827 LUAD cells expressed low FoxO1 but appreciable FoxO3a (**Supplementary Figure S5A**). Therefore, we focused on FoxO3a for subsequent investigations. Phosphorylation of FoxO3a at Ser294 and Ser425 leads to nuclear export of FoxO3a and cytosolic retention, resulting in loss of transcriptional activity and degradation[28]. Treatment of *EGFR*-mutant H1650 and HCC827 LUAD cells with gefitinib repressed phosphorylation of FoxO3a at Ser294 and Ser425 residues, reduced cytosolic FoxO3A and promoted nuclear translocation, suggesting transcriptional activation (**Figure 3A-B**). Similar results were found when treating respective cells with 5-FU or gemcitabine (**Supplementary Figure S6A-B**). Furthermore, overexpression of FoxO3a in both H1650 and HCC827 cell lines resulted in upregulation of basal ALPP expression, which was further potentiated when respective cancer cells were treated with gefitinib (**Figure 3C**). Inversely, inhibition of FoxO3A using the small molecule inhibitor AS1842856 [29, 30] diminished ALPP protein expression and blunted gefitinib-mediated ALPP upregulation in H1650 LUAD cells (**Figure 3D-E**).

**Figure 3.**
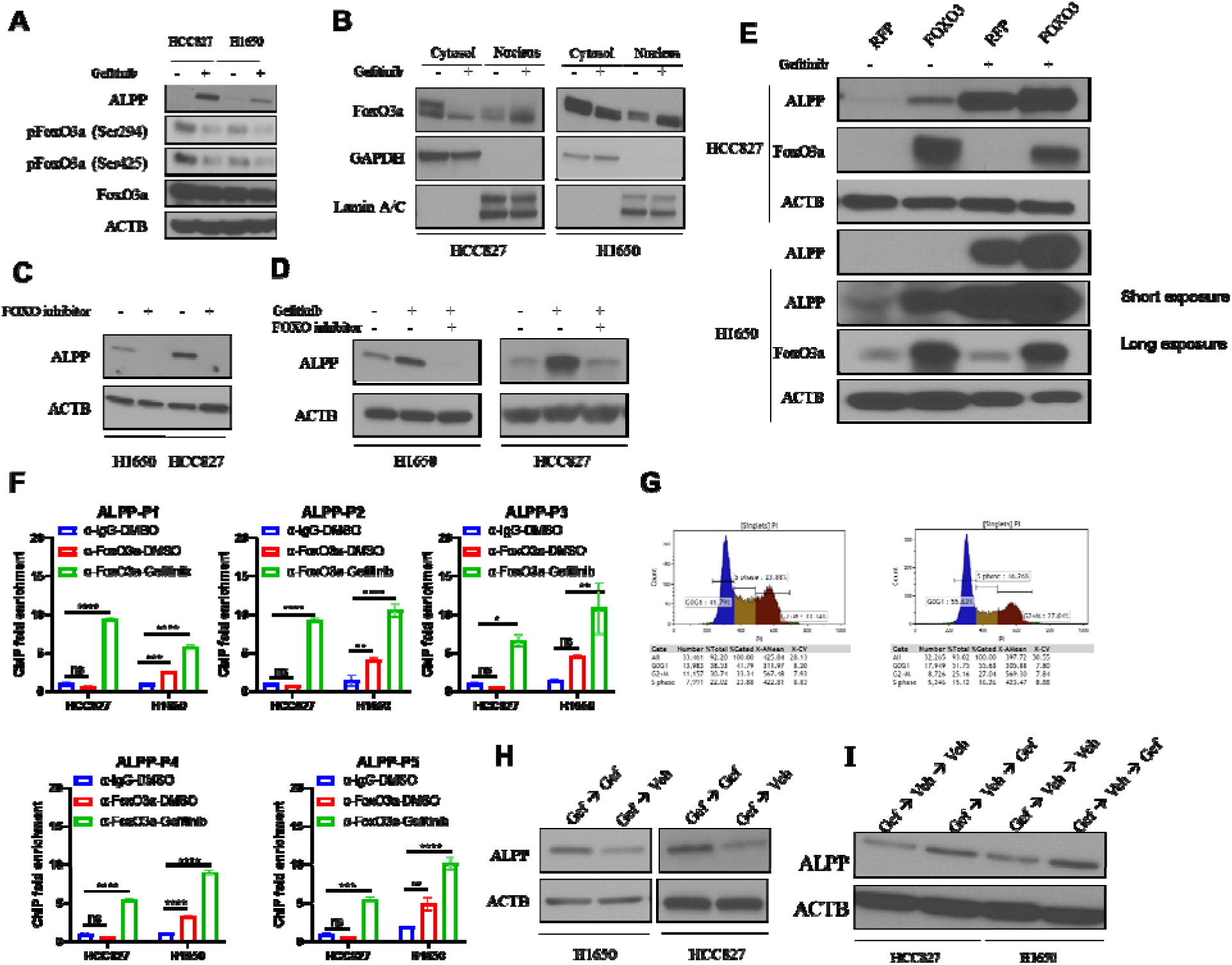
ALPP is transcriptionally regulated by FoxO3a and ALPP expression correlates with cancer cell quiescence. **(A)** Immunoblots for ALPP, FoxO3a and phosphorylated FoxO3a (Ser294, Ser425) in HCC827 and H1650 LUAD cells treated with 1 µM gefitinib for 6 hours. **(B)** Immunoblots for FoxO3a in the cytosol and nucleus compartment from gefitinib-treated HC827 and H1650 LUAD cells. **(C)** Immunoblots for ALPP in HCC827 and H1650 LUAD cells following 48-hour treatment with either vehicle or FoxO inhibitor AS1842856 (1 µM). **(D)** Immunoblots for ALPP in HCC827 and H1650 LUAD cells following 48-hour treatment with gefitinib (1 µM) plus either vehicle or FoxO inhibitor AS1842856 (1 µM). **(E)** Immunoblots for ALPP and FoxO3a in FoxO3a-overexpressing HCC827 and H1650 LUAD cells treated with either vehicle or gefitinib (1 µM) for 48 hours. **(F)** ChIP-qPCR assay for the promoter region of *FOXO3A* gene in HCC827 and H1650 LUAD cells treated with either vehicle or gefitinib (1 µM) for 6 hours. **(G)** Flow cytometry assay for PI-stained H1650 LUAD cells treated with vehicle or gefitinib (1µM) for 48 hours. **(H)** Immunoblots for ALPP in HCC827 and H1650 LUAD cells treated with gefitinib (1 µM) for 48 hours, followed by washing out of gefitinib, and treated with vehicle or gefitinib (1 µM) for another 48 hours. **(I)** Immunoblots for ALPP in HCC827 and H1650 LUAD cells treated with gefitinib (1 µM) for 48 hours, followed by washing out of gefitinib and rested for 48 hours. Cells were then treated with vehicle or gefitinib (1 µM) again for another 48 hours.

To elucidate whether FoxO3a was the transcriptional regulator of ALPP, we first performed in silico predictions using the online tool PROMO to predict the putative transcription factor(s) for ALPP and identified 8 potential binding sites for FoxO3a within 2 kb promoter region upstream of *ALPP* gene (**Supplementary Figure S5B**). We next performed ChIP-qPCR and confirmed that gefitinib treatment promoted FoxO3A binding to *ALPP* promoter sequences with resultant increases in ALPP mRNA levels in both H1650 and HCC827 LUAD cells (**Figure 3F-G**). Taken together, these data demonstrated that gefitinib suppresses EGFR signaling resulting in FoxO3A nuclear translocation and FoxO3A-mediated transcription of ALPP.

### ALPP expression correlates with cancer cell quiescence

High confluency contact inhibition, serum-deprivation as well as targeted agents, such as TKIs, can induce cellular quiescence [31]. FoxO3a is essential for maintaining cell quiescence [32, 33, 34]. Given our findings that FoxO3a transcriptionally regulates ALPP, we assessed whether ALPP associated with cancer cell quiescence. Flow cytometry analyses of propidium iodide stained H1650 cells revealed significant higher percentage of gefitinib-treated cells sequestered in G0/G1 phase (**Figure 3H**). To interrogate if the induction of ALPP is permanent or transient, H1650 and HCC827 LUAD cells were initially treated with gefitinib followed by continued culturing in gefitinib-containing medium or transition to medium containing vehicle (DMSO) control. ALPP levels were decreased in the absence of gefitinib (**Figure 3I**). Moreover, ALPP expression increased when LUAD cells that were transition to vehicle containing-medium were re-challenged with gefitinib (**Figure 3J**). Collectively, these data prove supportive evidence that ALPP expression is transient and associated with cancer cell quiescence status.

### Gefitinib treatment potentiates tumoral ALPP expression and enhances anti-cancer efficacy of ALPP-MAF-ADC *in vivo*

We targeted surface ALPP using Phrodo-Red conjugated ALPP antibody in H1650 LUAD cells and demonstrated internalization (**Figure 4A**), reinforcing utility of ALPP as a candidate for antibody-drug conjugate (ADC) therapy. We subsequently synthesized anti-ALPP antibodies conjugated with monomethylauristatin F (MMAF), the microtubule-disrupting agent inducing cell cycle arrest and apoptosis and tested the anti-cancer efficacy of gefitinib and ALPP-MMAF-ADC alone and the combination *in vitro* using H1650 LUAD cells. The combination treatment showed synergistic effects with potent cancer killing effects (**Figure 4B-C**). Similarly, findings were found in *EGFR*-wt H1651 and H2291 LUAD cells (**Figure 4D**).

**Figure 4.**
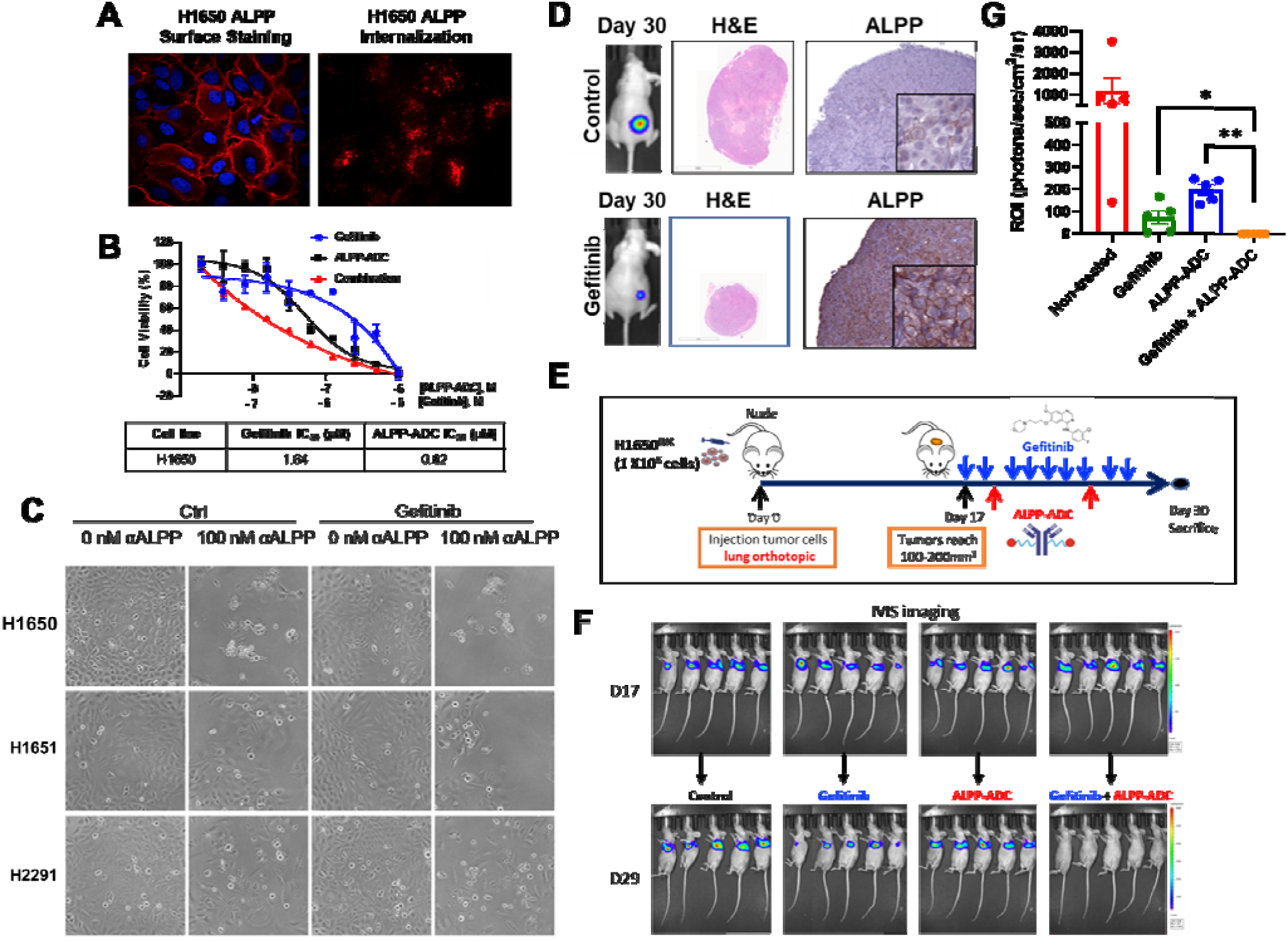
Gefitinib treatment potentiates tumoral ALPP expression and enhances anti-cancer efficacy of ALPP-MAF-ADC *in vivo*. **(A)** Confocal imaging of H1650 cells treated with or without Phrodo Red labelled anti-ALPP antibody. **(B)** Cell viability of H1650 cells treated with serial gefitinib and/or ALPP-MMAF-ADC. **(C)** Morphology of LUAD cell lines H1650, H1651, H2291, and H1944 treated with gefitinib and/or ALPP-MMAF-ADC. **(D)** H&E staining of tumor tisue and IHC staining of ALPP in tumor tissues from subcutaneous xenograft LUAD mouse model of H1650 cell line treated with or without gefitinib (50mg/kg). **(E)** Schematic of treating of orthotopic xenograft LUAD mouse model of H1650 cell line with gefitinib (50mg/kg) and/or ALPP-MMAF-ADC (5 mg/kg). **(F)** IVIS imaging of tumors before drug treatment at Day 17 and post-treatment at Day 29. **(G)** Statistical analysis of tumor burden from **(F)**. * *P*<0.05, ** *P*<0.01.

**Figure 5.**
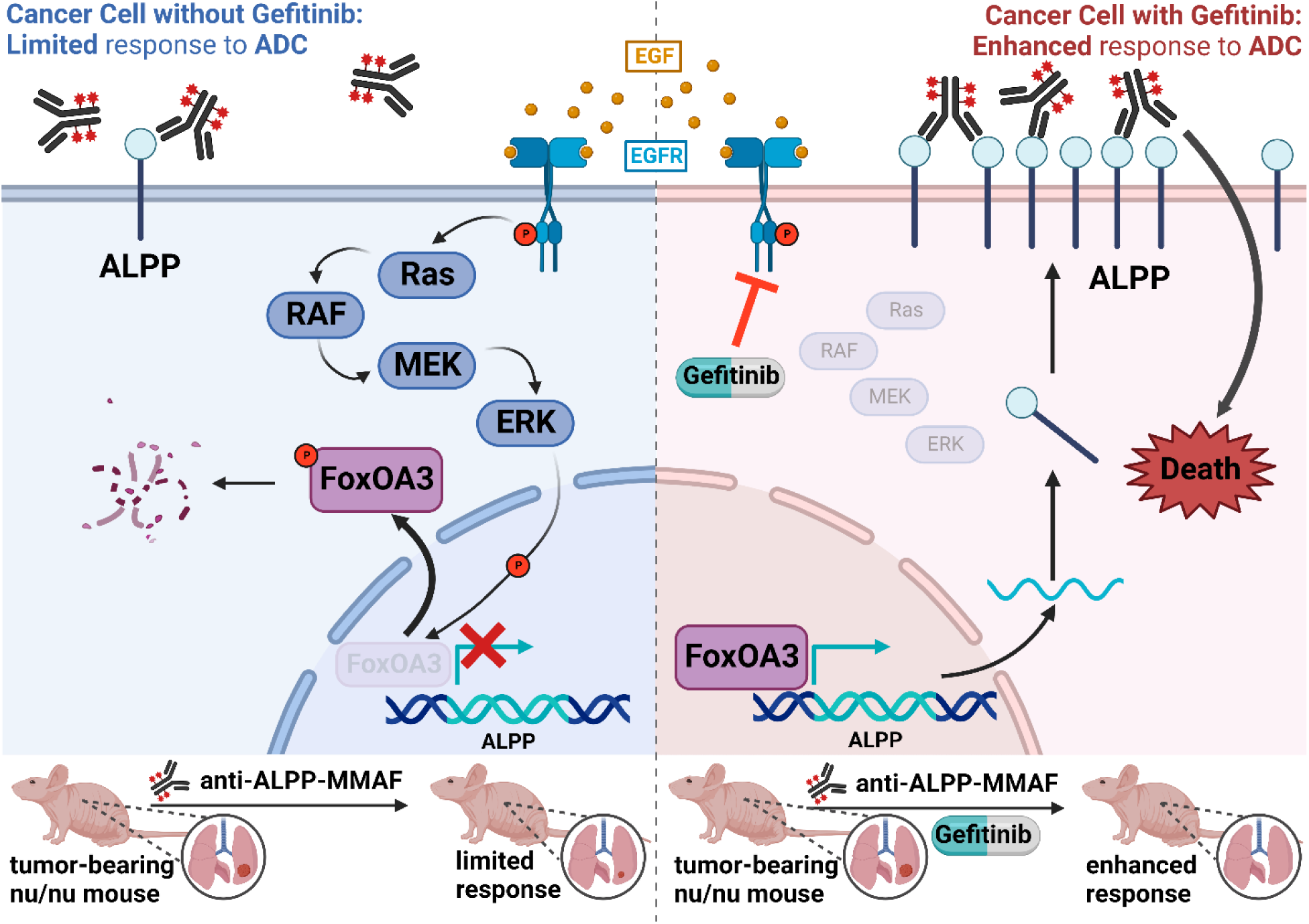
Proposed Schematic of ALPP surface upregulation in cancer cells by EGFR inhibition.

We next orthotopically implanted firefly luciferase-expressing H1650 (H1650^Luc^) LUAD cells into BALB/c nude mice and challenged tumor bearing mice daily with (50mg/kg) gefitinib or saline control for 10 days; tumor and other organ tissues were subsequently harvested and ALPP expression assessed via IHC. Consistent with our in vitro observations, tumor-bearing mice treated with gefitinib had pronounced increases in tumoral ALPP surface expression (**Figure 4E**). ALPP protein staining in other tissues were negative or showed no appreciable difference compared to vehicle control (**Supplemental Figure S7**). Using an independent cohort of H1650^Luc^-tumor bearing mice, we assessed the anti-cancer efficacy of 50mg/kg gefitinib, 5mg/kg ALPP-MMAF-ADC alone and in combination (**Figure 4F**). Treatment with gefitinib or ALPP-MMAF-ADC alone resulted in a statistically significant reduction in relative fluorescence units (RFU) of H1650^Luc^ tumors compared to control. Remarkably, tumor-bearing mice treated with gefitinib plus ALPP-MAFF-ADC had complete responses and were tumor free based on IVIS imaging (**Figure 4F-G**).

## Discussion

ALPP and ALPPL2 are closely related and regulated GPI anchored proteins that are known to be expressed on the cell surface in some cancers, whereas normal tissue expression is largely limited to the placenta [3, 6]. ALPP and ALPPL2 are currently being explored as cancer therapy targets [5, 7, 8, 9]. However, limited expression levels of ALPP/ALPPL2 in most cancer types may restrict its broader therapeutic potential. In the current study, we highlight a novel ‘two-hit’ therapeutic strategy of enhanced surface expression of ALPP in ALPP-expressing cancer cells, but not normal tissues, whereby treatment with therapeutics that target an oncogenic driver, e.g. EGFR, increases the efficacy of ALPP-targeting therapies.

Consistent with prior studies [6], our assessment of ALPP and ALPPL2 mRNA expression and surface protein expression in human tumors and cancer cell lines demonstrated variable expression with outliers being frequently observed in lung adenocarcinomas as well as gastrointestinal and gynecological malignancies. In the TMA of lung tumors, ALPP surface staining was found in 11.3% of all lung tumors analyzed, with the highest frequency of ALPP surface staining being observed in LUAD (17.6%). Notably, ALPP staining positivity was associated with smoking status. Prior studies have reported that serum levels of ALPP and ALPPL2 are increased up to 10-fold in individuals that smoke cigarettes compared to non-smokers.[35, 36] Moreover, epigenome-wide association study (EWAs) comparing current, former and never smokers from 1,793 participants revealed that methylation of CpG islands in the *ALPP/ALPPL2* loci for smokers was lower than that of non-smokers.[37]

Prior studies have documented an inverse relationship between ALPP expression and cell proliferative status [19, 20, 21, 22, 23]. Consistent with these observations, we found marked ALPP upregulation in cancer cells that were subjected to growth-inhibitory stimuli including serum restriction, cell-cell contact inhibition, or treated with anti-proliferative chemotherapeutic 5-fluorouracil or gemcitabine or EGFR-targeting tyrosine kinase inhibitors. Of importance, enhancement of ALPP expression in cancer cells was transient and reversible, suggesting that ALPP is may be associated with cell quiescence [32, 38, 39, 40, 41, 42, 43, 44]. In support of this, we identified FoxO3a, a key regulator of cell quiescence [32, 33, 34], as the upstream transcriptional regulator of ALPP. FoxO3a is one of four related FoxO transcription factors that protect cells against a wide range of physiologic stresses, playing central roles in DNA repair, growth arrest, and apoptosis in response to DNA damage and oxidative stress.[45] FoxO3a inactivation is frequently reported in various malignancies. Inactivation of FoxO3a in tumors has been attributed to overactivation of the PI3K-AKT or MEK-ERK signaling pathways, which negatively regulate FoxO3a activity via phosphorylation of FoxO3a at Thr32, Ser253 and Ser294 that prompts and sustains nuclear exclusion to the cytoplasm and proteasomal degradation [28, 46, 47, 48]. In our study, we found that gefitinib treatment suppresses FoxO3a phosphorylation in LUAD cells, resulting in FoxO3a nuclear translocation that was met with concomitant increases in ALPP mRNA and protein levels and elevated surface ALPP expression. Inhibition of FoxO3a using the small molecule inhibitor AS1842856 [29, 30] reduced ALPP protein expression and prevented gefitinib-mediated ALPP upregulation. Moreover, treatment of LUAD cells with anti-proliferative chemotherapeutic agents 5-FU and gemcitabine similarly promoted FoxO3a dephosphorylation and nuclear translocation, and increased ALPP protein levels, thus providing a converging mechanism of FoxO3a-mediated ALPP upregulation in cancer cells.

Our findings prompted us to evaluate whether gefitinib-mediated enhancement surface ALPP expression in cancer cells may increase the efficacy of ALPP-targeting therapies. To this end, we demonstrated that combination treatment of LUAD cells with gefitinib plus ALPP-ADC-MAF resulted in markedly improved anti-cancer killing effects compared to either treatment modality alone. Treatment of *EGFRmutant* LUAD-tumor bearing mice with gefitinib similarly enriched ALPP surface expression in tumors but not in normal tissues. Combinatorial treatment of *EGFRmutant* LUAD-tumor bearing mice with gefitinib plus ALPP-ADC-MAF resulted in pronounced anti-cancer effects compared to gefitinib or ALPP-ADC-MAF treatment alone. Although gefitinib is an older generation TKI, our *in vitro* studies demonstrated that other TKIs, including lapatinib, afatinib, and osimertinib, similarly enhanced ALPP expression in LUAD cells. Thus, we believe that newer generation TKIs would yield similar results *in vivo*.

In conclusion, our study provides mechanistic insights into ALPP upregulation in cancer cells and establishes FoxO3a as the transcriptional regulator of ALPP. Importantly, we provide a novel combinational therapeutic strategy of enhanced ALPP expression in cancer cells that has potential to increase efficacy of ALPP-targeting therapies.

## Supplemental Figures and Tables

**Supplemental Table S3.** ALPP Membrane Staining Positivity in Tissue Microarray of Lung Adenocarcinoma and Squamous Cell Carcinoma.

| Variable | Membrane Staining |  | P-value† |
| --- | --- | --- | --- |
|  | ALPP Positive | ALPP Negative |  |
| Total | 23 | 181 |  |
| Sex, N (%) |  |  |  |
| Male | 8 (34.8) | 98 (54.1) | 0.12 |
| Female | 15 (65.2) | 83 (45.9) |  |
| Smoking Status, N (%) |  |  |  |
| Current | 14 (60.9) | 49 (27.1) | 0.0001 |
| Former | 7 (30.4) | 51 (28.2) |  |
| Never | 0 (0) | 17 (9.4) |  |
| Unknown | 2 (8.7) | 64 (35.4) |  |
| Subtype, N (%) |  |  |  |
| Adenocarcinoma | 21 (91.3) | 119 (65.7) | 0.015 |
| Squamous Cell Carcinoma | 2 (8.7) | 62 (34.3) |  |
| Stage, N (%) |  |  |  |
| I+II | 15 (65.2) | 125 (69.1) | 0.81 |
| III+IV | 8 (34.8) | 56 (30.9) |  |
| Mutation Status, N (%) |  |  |  |
| KRASmut | 5 (21.7) | 32 (17.7) | 0.45 |
| EGFRmut | 1 (4.3) | 15 (8.3) |  |
| KRASwt/EGFRwt | 13 (56.5) | 59 (32.6) |  |
| Unknown | 2 (8.7) | 75 (41.4) |  |
| Neoadjuvant Therapy, N (%) |  |  |  |
| Yes | 1 (4.3) | 32 (17.7) | 0.10 |
| No | 22 (95.7) | 148 (81.8) |  |

† Statistical significance was determined by Fisher's Exact Test for two class categorical comparisons or $\chi^2$ test for trend for multiple categorical variables. 2-sided P-values are reported. Abbreviations: wt- wildtype

**Supplemental Figure S1.**
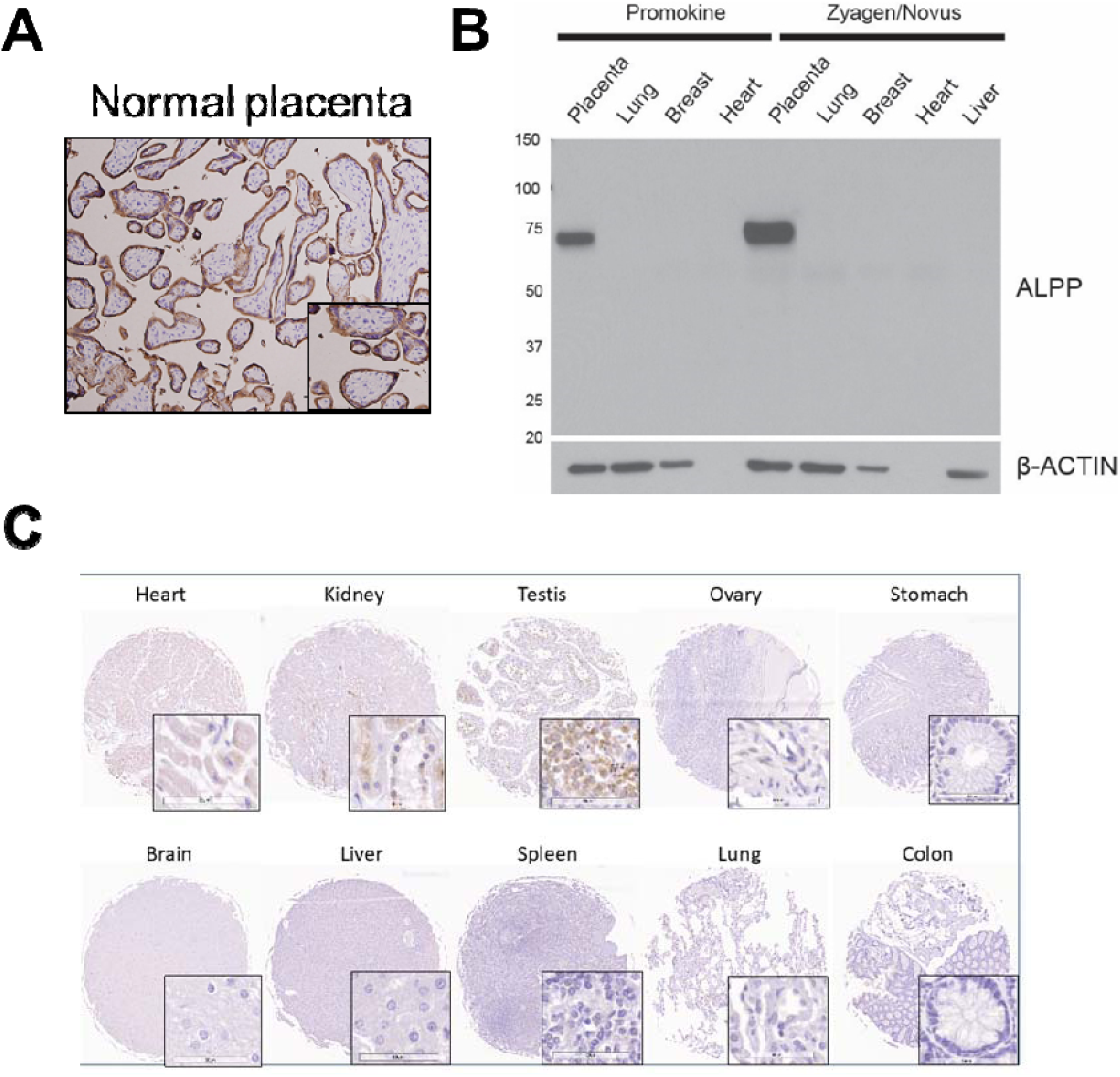
ALPP expression in normal tissues. **A)** Representative immunohistochemistry (IHC) image for ALPP staining in placental tissues. **B)** immunoblots for ALPP in protein lysates from placenta, lung, breast, and heart tissues. **C)** Representative IHC sections for ALPP in various normal tissues.

**Supplemental Figure S2.**
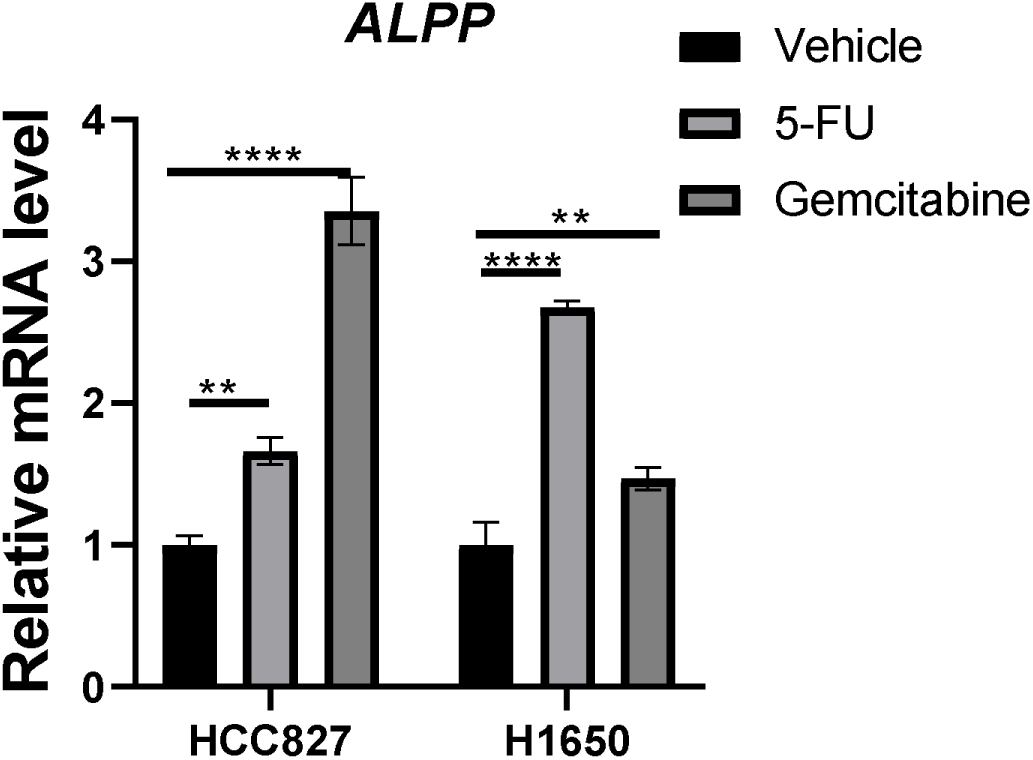
5-FU and gemcitabine potentiate ALPP in LUAD cell lines. qPCR for *ALPP* in EGFR-mutant H1650 and HCC827 LUAD cells following 24hour treatment with vehicle control (DMSO), 5-FU (10 µM) or gemcitabine (10 nM).

**Supplemental Figure S3.**
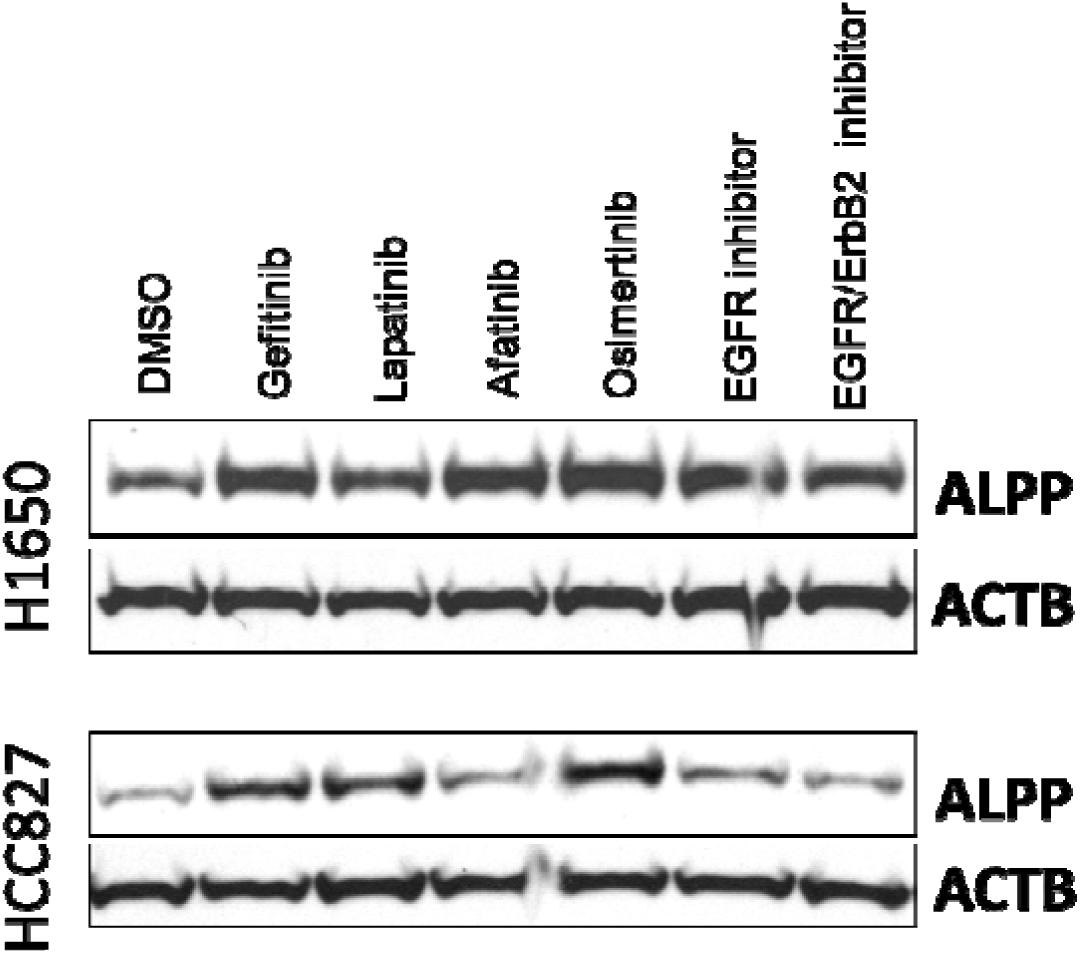
EGFR-targeting TKIs potentiate ALPP in LUAD cell lines. **Immunoblots for ALPP and ACTB (loading control) in EGFR-mutant** H1650 and HCC827 LUAD cells following 24hour treatment with vehicle control (DMSO), or 1uM of varying TKIs.

**Supplemental Figure S4.**
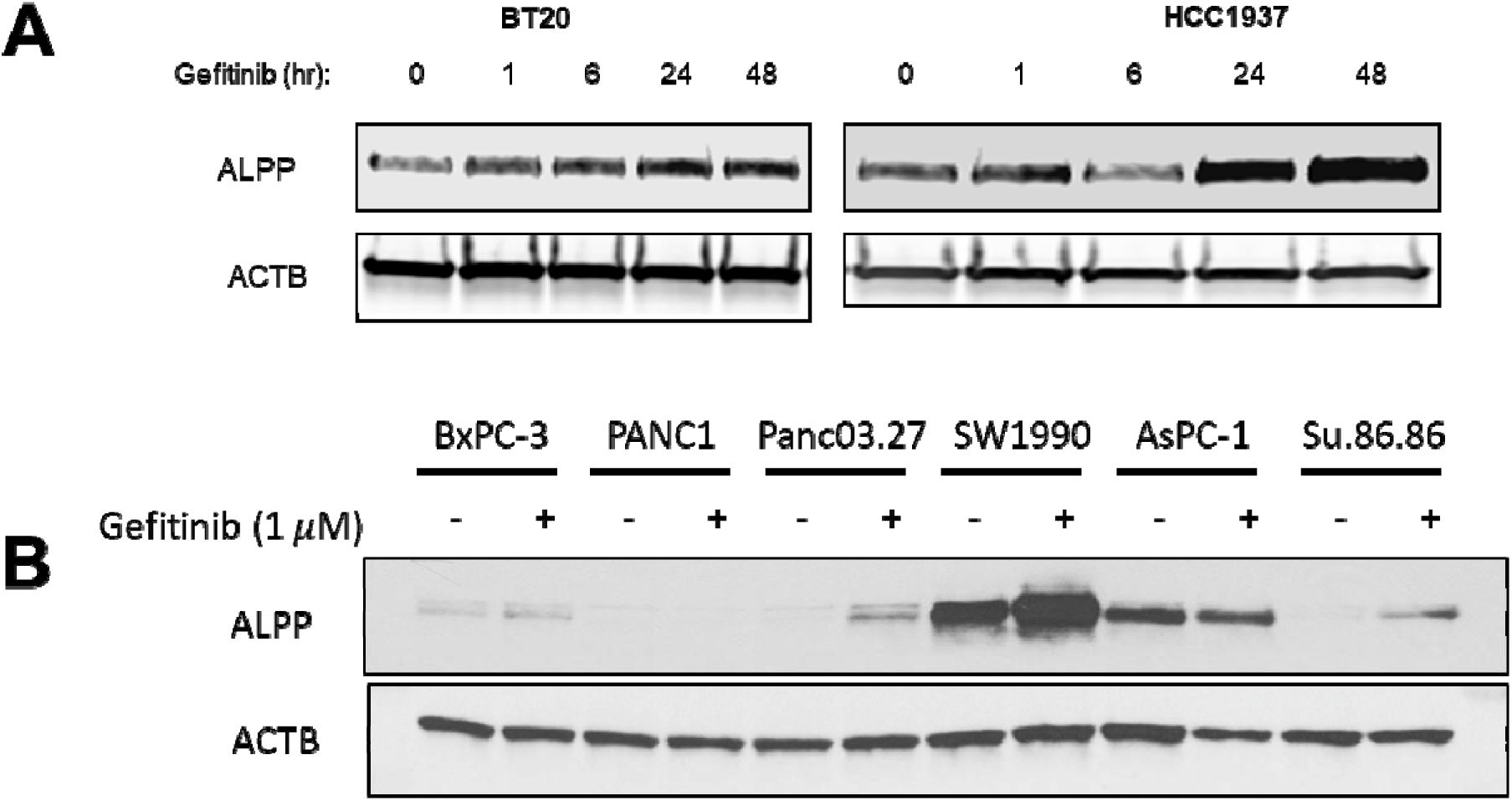
EGFR inhibition with gefitinib potentiates ALPP expression in other cancer types. Immunoblots for ALPP and ACTB (loading control) in triple-negative breast cancer cell lines BT20 and HCC1937 (A) or pancreatic cancer cell lines (B) following treatment with 1μM gefitinib. For panel B, pancreatic cancer cell lines were treated for 24 hours with either vehicle (DMSO) or 1μM gefitinib.

**Supplemental Figure S5.**
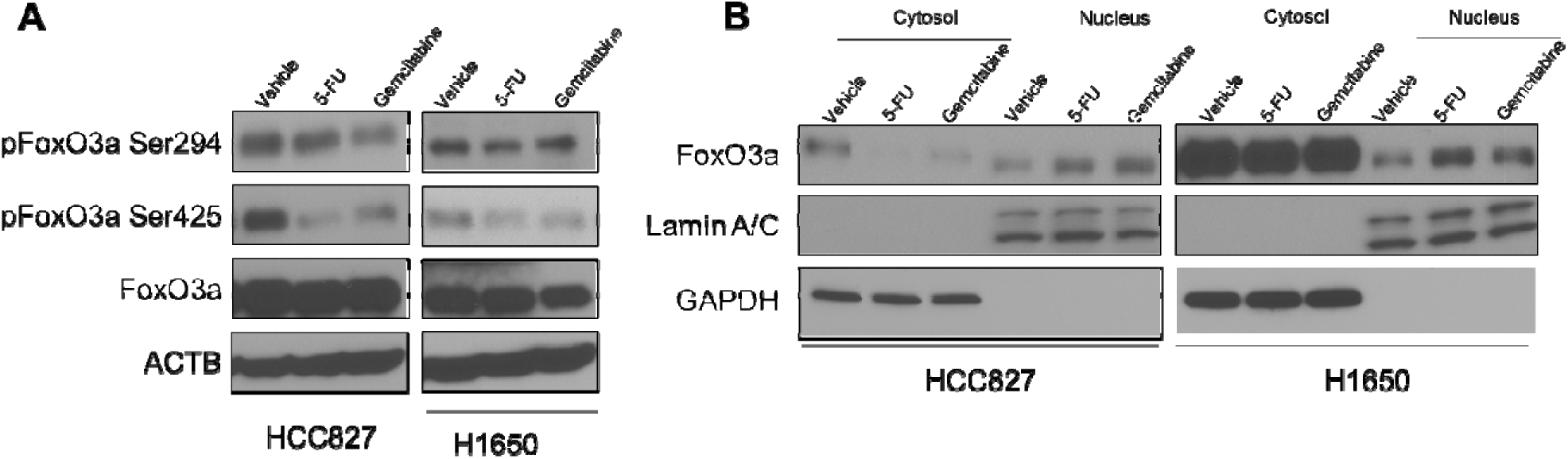
5-FU and gemcitabine promote FoxO3a activation. **(A)** Immunoblots for ALPP, FoxO3a and phosphorylated FoxO3a (Ser294, Ser425) in HCC827 and H1650 LUAD cells treated with vehicle, 5-FU (10 µM) or Gemcitabine (10 nM) for 6 hours. **(B)** Immunoblots for FoxO3a in the cytosol and nucleus compartment from vehicle, 5-FU (10 µM) or Gemcitabine (10 nM)-treated HC827 and H1650 LUAD cells.

**Supplemental Figure S6.**
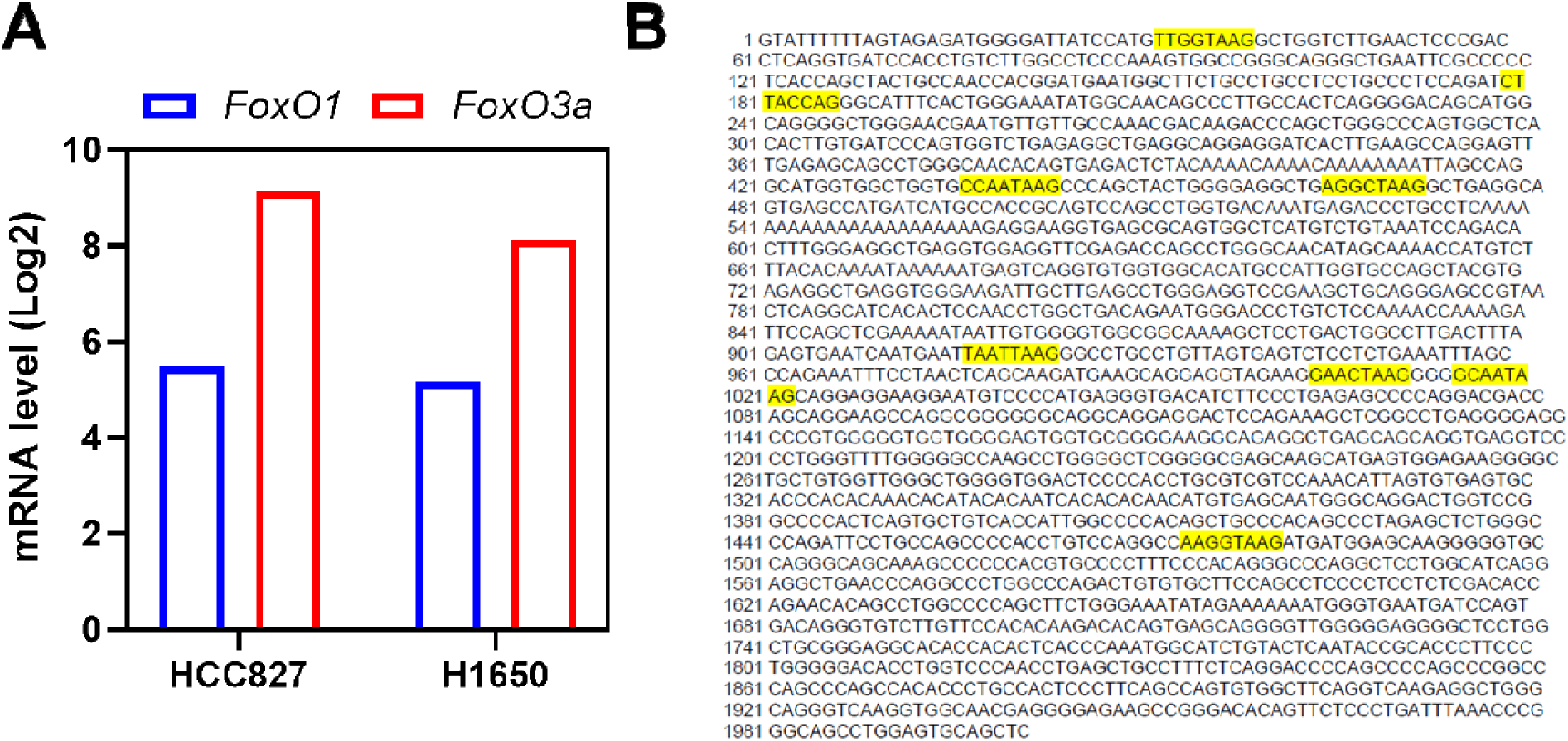
Potential FoxO3a binding sites in the promoter region of *ALPP* gene. **A)** mRNA expression of FoxO1 and FoxO3a in HCC827 and H1650 LUAD cells. Data was derived from The Cancer Cell Encyclopedia (CCLE) database. **B)** Putative FoxO3a binding sites in the promotor region of ALPP.

**Supplemental Figure S7.**
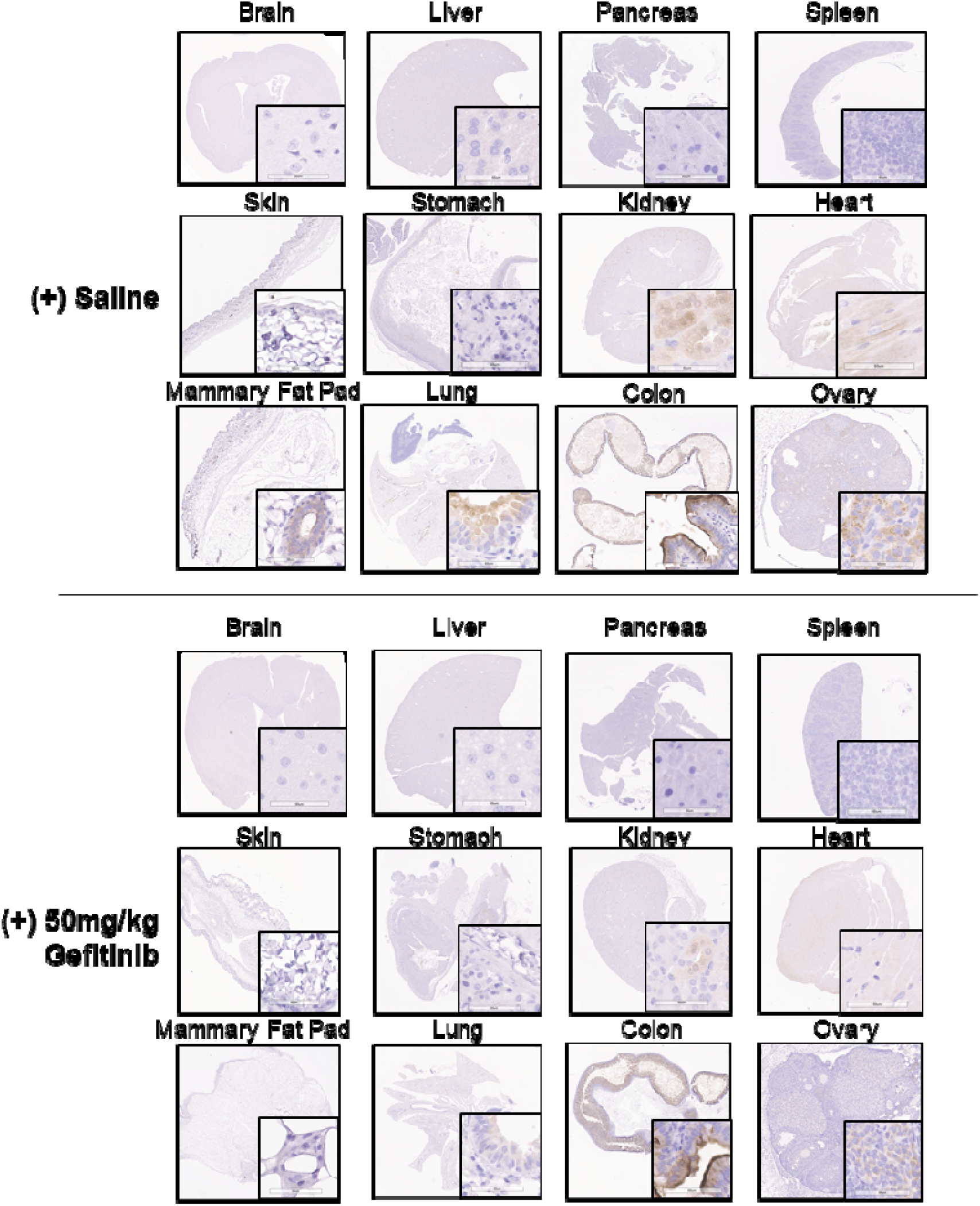
IHC analyses for ALPP in normal tissues of tumor bearing mice following gefitinib treatment.

